# Neural Dynamics of Belief and Value Computations Guiding Strategic Social Decisions

**DOI:** 10.64898/2026.05.08.723744

**Authors:** Arkady Konovalov, Jie Hu, Christian C. Ruff

## Abstract

Successful strategic behavior must be grounded in beliefs about the opponent and her intentions. While many potential models have been proposed to explain choices in such situations, the neural mechanisms that govern learning and choice in complex strategic contexts remain poorly understood. Here, we use a computational model that combines dynamic learning and choice mechanisms to explain both choices and response times of human participants engaged in a competitive strategic task. Using electroencephalography (EEG), we identify temporally structured stages of neural processing that support an evolving value-based decision process, corresponding to first- and second-order belief updates and evidence accumulation related to the comparison of action values. Gamma-band phase coupling between central and parietal EEG signals varied with individual winning rate, suggesting that strategic behavior involves coordinated information transfer across spatially remote areas. Together, our data characterize the temporally evolving neural dynamics of belief and valuation processes that underlie strategic choice and provide neural validation for assumptions embedded in computational models of this behavior.

## INTRODUCTION

Repeated strategic interactions are common in everyday life, including sports, market competition, and politics. These interactions often require learning the distribution of the opponent’s past actions and understanding the consequences of one’s own past actions to predict future moves and determine the best behavioral strategy (Camerer, 2003). In game theory and behavioral science, such processes are well explained by models of dynamic learning and belief updating (Camerer & Ho, 1999; Hampton et al., 2008; Roth & Erev, 1995). These models usually rely on assumptions about the hidden processes behind observed choices. Specifically, they involve different processes believed to perform parallel or sequential calculations of hidden variables: Strategic decisions have values (or utilities, or “attractions”) computed from potential outcomes (typically payoffs to the players) and the players’ beliefs. As the interaction evolves, these beliefs are dynamically updated using the history of play. Individuals can form first-order beliefs about the opponent’s actions (often called “fictitious play” (Boylan & El-Gamal, 1993)) and second-order beliefs, which involve mentalizing the opponent’s strategy, often represented through influence learning or level-k models (Buergi et al., 2026; Coricelli & Nagel, 2009; Hampton et al., 2008; Hill et al., 2017; Konovalov et al., 2021; Nagel, 1995). While experimental and market data support the predictive power of these models (Camerer, 2003), without neural data, it is often unclear whether belief-updating models only approximate the brain’s decision-making mechanisms or actually reflect the underlying computational processes.

For this reason, various studies have empirically validated some assumptions of strategic learning models through functional magnetic resonance imaging (fMRI) experiments, showing that the trial-by-trial values and beliefs embedded in these models vary dynamically and correlate with activity in different brain regions (Bhatt et al., 2010; Camerer & Hare, 2014; Hampton et al., 2008; Hill et al., 2017; Konovalov, 2026; Konovalov et al., 2018, 2021; Konovalov & Ruff, 2022; Park et al., 2019; Seo et al., 2014). Specifically, reward prediction errors in strategic contexts were associated with activity in the striatum (Set et al., 2014; Zhu et al., 2012), first-order beliefs were linked to activity in the dorsomedial prefrontal cortex (dmPFC) (Hampton et al., 2008; Set et al., 2014; Zhu et al., 2012), and second-order beliefs were causally connected to activity in the right temporoparietal junction (rTPJ) (Hampton et al., 2008; Hill et al., 2017). Further evidence indicates that the dmPFC may be involved in general strategy implementation, while the TPJ might provide immediate updates to the decision variables (Konovalov et al., 2021).

While some of these regions are part of a network often called the “social brain” (Adolphs, 2009; Frith, 2007; Lockwood et al., 2020), many of these computations may not be explicitly driven by social context but instead reflect the reactive nature of a strategic environment (Konovalov et al., 2021). Importantly, all these models assume that each individual decision is ultimately guided by the subjective values of actions, which incorporate dynamically changing beliefs about the opponent’s actions and the effects of one’s own actions on the opponent’s play. These action values have often been linked to activity in the ventromedial prefrontal cortex (vmPFC) (Bartra et al., 2013; Clithero & Rangel, 2014); however, the precise process by which these values are converted into actual actions remains poorly understood.

While fMRI can map the topographical organization of these updates in the brain, it lacks temporal resolution, and many questions remain. Do first- and second-order belief processing and value comparison occur simultaneously or sequentially? Do they coincide with reward processing? Are decision values computed and compared during belief updating or at the time of choice? How are learned decision values transformed into actions? Here, we explore these questions by leveraging the temporal resolution of electroencephalography (EEG).

We aimed to integrate strategic learning (using belief-updating models) and value-based decision-making (using evidence-accumulation, or EA, models) within a single framework by modeling a dynamic readout of the corresponding neural signals that evolve at millisecond resolution during a trial. This approach differs significantly from fMRI, which provides only static snapshots of processes occurring at clearly distinct temporal stages of a trial, and only if these stages are separated by several seconds. Our method enabled us to investigate model-based learning and decision-making at millisecond resolution in a complex strategic environment in which the values of identical actions vary across trials. This reflects the dynamic nature of latent beliefs and decision-making processes, rather than the static subjective values that have been investigated in many other EEG studies of, e.g., value-based choices (Hu et al., 2023; Pisauro et al., 2017; Polanía et al., 2014) or perceptual decision tasks (Kelly & O’Connell, 2013; O’Connell et al., 2012; Ratcliff et al., 2009).

Our approach incorporates the principles of evidence-accumulation models, which posit that the brain performs a dynamically evolving comparison of values to choose the more advantageous action (Lin et al., 2020; Liu & Pleskac, 2011; Polanía et al., 2014; Ratcliff & McKoon, 2008). These models can explain choices and response times (RT) across various tasks, including perceptual, memory, and economic decisions (Bogacz et al., 2010; Konovalov & Krajbich, 2019; Krajbich et al., 2012; Pedersen et al., 2017; Polanía et al., 2014). Neural studies using EEG and single-neuron recordings have shown that some ramp-up signals observed during decision-making may be explained by evidence-accumulation models; however, most of these studies focus on simple perceptual and preference-based decisions. Only one recent study has shown that evidence accumulation can be detected during non-interactive social decision-making using EEG (Hu et al., 2023), but decisions in this context reflected static social preferences rather than dynamically changing beliefs and action values, as required in our competitive strategic setting.

We combined the dynamic learning and evidence-accumulation methods and adapted a commonly used Inspection game for the EEG setup. In this strategic game, one player (in our case, a participant in the EEG room) played a simple two-action game against another human opponent in a different room (Figure 1A) (Hampton et al., 2008; Hill et al., 2017). Compared to previous versions, we simplified the visual presentation to reduce noise in the EEG signal: The EEG participant chose between two options, labeled H and T, and then observed whether the opponent selected UP or DOWN (Figure 1B). The player (and the opponent) could win in two of four possible action combinations, with these combinations being complementary and the opponents’ goals thus mismatched. Participants learned the entire outcome space before the game, including their own and the opponent’s payoffs, and based their decisions on the history of play. The players completed 400 trials, divided into eight blocks.

**Figure 1.**
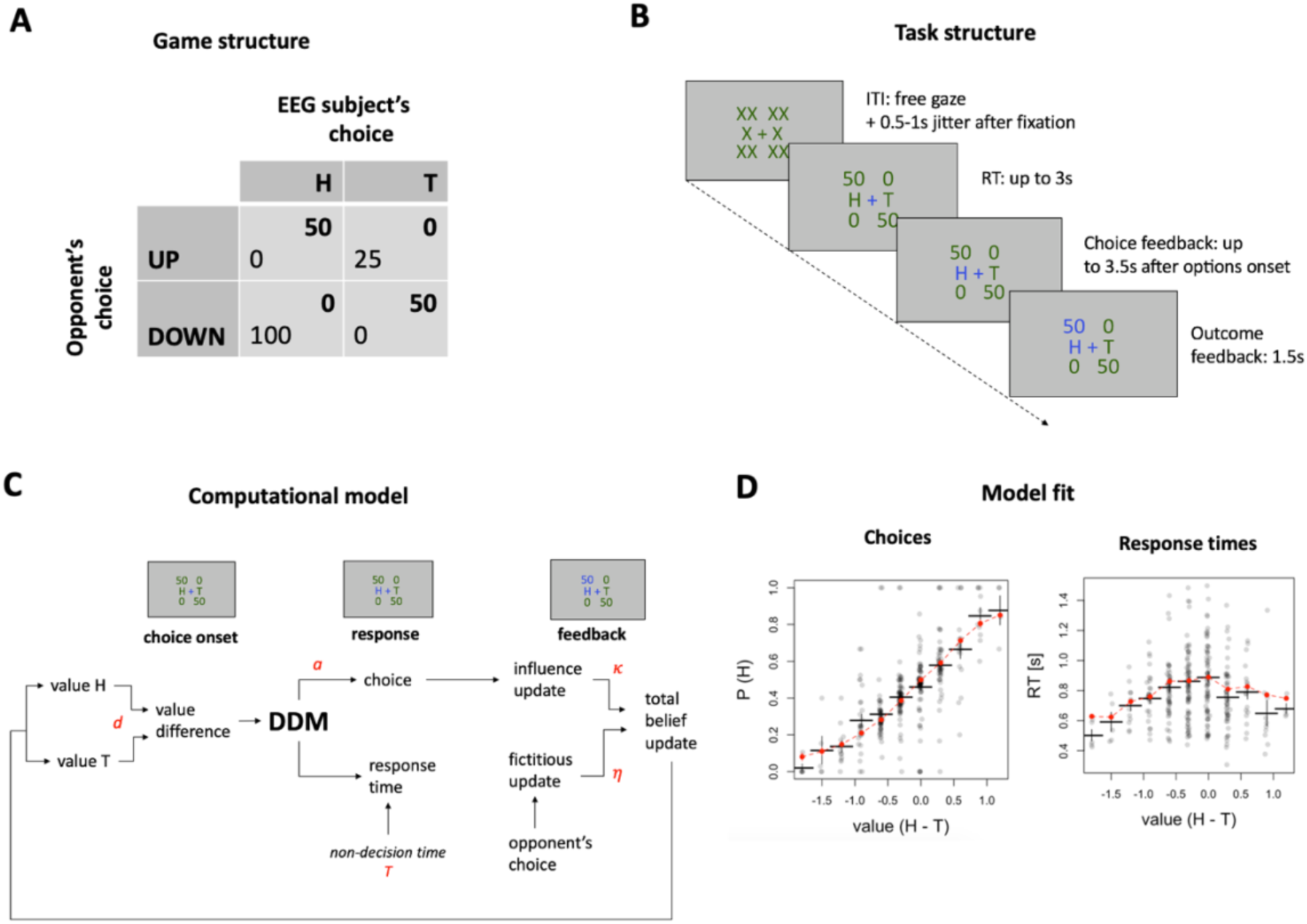
(A) Game matrix. The EEG participant chose between H and T (columns), with the opponent’s choices labeled as UP and DOWN (rows). The table shows the payoffs for both participants based on their choices (the EEG participant’s payoffs are in bold). (B) Task structure. Before the trial began, the choice options and payoffs were hidden with “X” letters. The participant was required to fixate on the central cross to continue (which then turned blue). The trial started after a randomly jittered interval of 0.5-1 seconds. The options and payoffs were then revealed (with H and T placements randomized each trial). The participant had 3 seconds to respond. The selected option changed color to blue; feedback was shown at 3.5 seconds after the trial began. The outcome (the opponent’s choice) was then displayed for 1.5 seconds, with the participant’s payoff highlighted in blue. (C) Computational model. The model assumed that the participant assigned dynamic values to options H and T. The value difference in each trial served as input in a DDM choice model. The choices made by the participant and opponent updated the participant’s beliefs about the opponent’s next choice. The overall belief influenced the future values of the options used for the next decision. The participant-specific free parameters are marked in red. See Methods for details. (D) Model fits and posterior predictive checks. The left panel shows the probability of choosing H based on the difference in model-generated values of H and T (the red line depicts the posterior predictive curve from model simulations). The right panel presents response times (RT) in seconds as a function of the value difference between H and T (with red indicating model simulation). Each dot represents an individual participant in the respective bin, and error bars denote standard errors of the mean (s.e.m.).

While standard game theory predicts that players should randomize between two actions (playing the so-called mixed-strategy equilibrium), in practice, humans find this very difficult (Camerer, 2003). The optimal strategy when facing an opponent who does not randomize is to track the opponent’s behavior and respond accordingly, as real human opponents tend to show autocorrelations in their choice history: most pay attention to their own and their opponent’s past play and make decisions based on that information (Camerer, 2003; Camerer & Hare, 2014; Hampton et al., 2008). To account for how option values change dynamically based on this history, we used a computational model that includes tracking the participant’s beliefs through an influence learning process (Hampton et al., 2008; Hill et al., 2017) and a value-comparison process using the standard drift-diffusion model (Ratcliff, 1978). We used a 128-channel EEG system sampling cortical activity at 500 Hz, which allowed us to compute neural correlates of trial-level variables such as belief updates and choice values, and to analyze evidence accumulation within trials using a model that combines learning with DDM-based decision-making.

To preview the results, we found strong support for the computational model integrating learning and evidence-accumulation processes. Our findings reveal a sequential decision process involving value comparison and belief updating, with first-and second-order beliefs processed during feedback and value comparison taking place during the choice stage. These results provide new evidence for several cognitive processes assumed by the model involved in strategic social decisions, offer insights into their temporal development, and suggest direct links between individual and social models of value-based decision-making.

## RESULTS

### The learning-DDM captures the identity and speed of strategic choices

To capture the parallel processes of social learning and decision-making, we used a simple, robust Inspection game paradigm that has been previously used to identify causal neural correlates of first- and second-order beliefs in the dmPFC and TPJ (Hampton et al., 2008; Hill et al., 2017). As described above, on each trial, the participant chose between two fixed options, H and T (Figure 1A). After making their own choice, they receive feedback about the opponent’s choice. We focused on each stage separately and identified the computations associated with the choice and feedback stage (Figure 1C). Crucially, the positions of options H and T were randomized on each trial, so participants had to wait for the choice screen to prepare and execute their response, ensuring that these motor-related processes could not confound our results.

To identify the value signals and beliefs of interest for the EEG analysis, we employed an extension of a model that was already used successfully in prior fMRI studies of this paradigm (Hampton et al., 2008; Hill et al., 2017), which we adapted by a sequential sampling-based choice model to yield predictions about dynamic evidence accumulation processes underlying strategic choices. Our model assumed that the participant formed subjective values of the H and T options and used these values to identify the best choice on each trial. To leverage EEG’s temporal precision, we did not use the standard softmax choice function typically employed in previous research, but instead used a drift-diffusion choice model (DDM) (Ratcliff & McKoon, 2008). This allowed us to fit response times (RT) in addition to choices and make predictions about evidence accumulation processes in the brain. Since our learning model included two parameters, we reduced computational complexity by employing a straightforward, robust version of the DDM (Ratcliff, 1978) that incorporates only three additional participant-level parameters: a decision boundary, a drift-rate multiplier, and a non-decision time.

While DDMs usually assume that choices on each trial are independent, more recent reinforcement learning–DDM (RL-DDM) approaches have been proposed for simple RL tasks (Fontanesi, Gluth, et al., 2019) where option values change on each trial. We adopted this approach here, assuming that the differences in the values of the options (H and T)—constructed based on beliefs about the opponent’s future actions, see below—enter the DDM as a drift rate (multiplied by a factor). We also considered an alternative model in which the value difference affected the starting-point bias, but this variant fit the data less well (see Methods).

To model learning, we assumed that, after receiving feedback on the opponent’s action, participants dynamically update their beliefs on each trial by using both the opponent’s actions and their own (Hampton et al., 2008; see Methods for a detailed description). Regarding first-order beliefs, participants monitor the relative frequency of the opponent’s choices (UP or DOWN), predict the opponent’s most likely next move, and respond by assigning value to the corresponding option, H or T, that would win against this predicted opponent action. Regarding second-order beliefs, participants track the relative frequency of their own choices and assume that this influences the opponent’s beliefs, which the opponent can also track as a first-order belief (see Methods for model details).

Following the previous literature (Hampton et al., 2008; Hill et al., 2017), we label the difference between the opponent’s choice on the current trial and their estimated frequency as the “fictitious play update”, and the weight of this update in the total belief update as a free participant-level parameter 𝜂 (see Methods). The difference between the participant’s own choice and the second-order belief (the probability calculated as an inverse choice function of the relative frequency of the participant’s own choices) on the current trial is the “influence update,” with the weight of this update in the total belief update designated as a free participant-level parameter 𝜅 (Figure 1B, Methods). We fit the model to each participant’s data using MLE (see Methods).

Posterior predictive checks confirmed that the model captured systematic variation in choices and RTs (Figure 1D), given the type of data and model class (see also (Konovalov et al., 2021)). It accurately classified 64% of choices (SD = 9%) per participant on average (in-sample, balanced prediction accuracy), aligning with previous findings for the influence learning model (Hill et al., 2017). RTs showed a typical DDM-predicted inverted-U relationship with the difference in subjective values, with easier decisions (i.e., those with a significant difference in subjective values) being faster (Figure 1D). Overall, these fits demonstrate that the relationship between choices and RT in our task aligns with a latent evidence-accumulation process, supporting our model’s assumptions about the dynamic link between values and choices during the decision phase. Note that in terms of perceptual information, the participant does *not* need to gather new information about the choice options on each trial (as is common in other perceptual and value-based evidence accumulation tasks (Frömer et al., 2024)), except for the left/right position of the response options. Therefore, evidence accumulation processes in our setting predominantly involve value information derived from beliefs about opponent actions.

To validate the sequential processes of value updating and to identify the dynamic neural correlates of the latent decision processes, we then used the model fits to compute several trial-by-trial model-based variables (value of the chosen option, value difference, fictitious and influence play updates; see Methods for the mathematical definitions) and inspect the EEG data for the corresponding signals.

### Temporally distinct processing of reward, first- and second-order belief updates

Previous fMRI research has identified the neural correlates of two key learning updates: first-order (fictitious play) and second-order (influence) updates, located in the dmPFC and TPJ, respectively (Hampton et al., 2008; Hill et al., 2017). A transcranial magnetic stimulation (TMS) study showed that disrupting activity in the right TPJ decreases the use of influence strategies in the Inspection game (Hill et al., 2017). Using our EEG data, we could now directly assess whether these two updates occur simultaneously or sequentially during the feedback stage, and whether they are encoded only during feedback or also during the choice stage. Furthermore, another important (explicit) event during feedback that we analyzed—separate from the belief updates—was the indication of reward outcome based on the choice made (0 vs 50).

When first examining the data for neural signatures of reward outcomes, we identified an amplitude correlate of reward occurring in the roughly 200-400 ms window after feedback onset, with positively correlated electrodes mainly clustering around the central area and negative activation concentrating in the parietal region (Figure 2A). The timing and topography of this activation align with previous literature (Martín & René, 2012; Meadows et al., 2016; Yeung & Sanfey, 2004). Importantly, in simultaneous EEG-fMRI studies, this central cluster of activity has been linked to activations in the vmPFC and nucleus accumbens, areas commonly associated with reward reception (Cohen et al., 2007; Flores et al., 2015; Gorka et al., 2015; HajiHosseini & Holroyd, 2015; Hoy et al., 2021; Keren et al., 2018; Silvetti et al., 2014).

**Figure 2.**
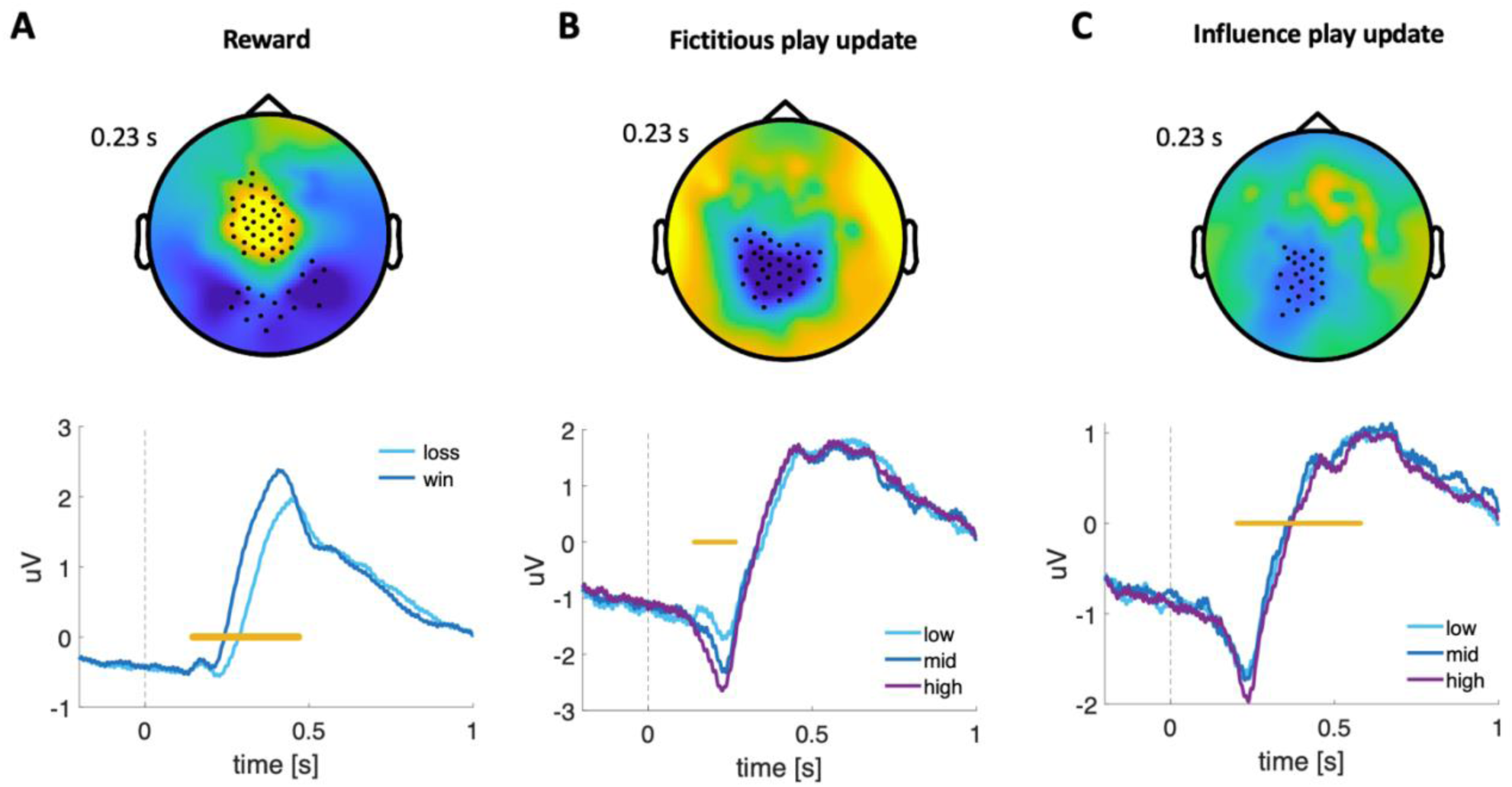
Feedback stage EEG results. The topographical maps show the significant clusters (marked with black dots) at 230 ms after feedback onset (timepoint 0). The yellow lines in the ERPs indicate the (approximate) time of significance at p < 0.05. (A) ERP of the EEG signal was calculated for significant electrode clusters during the feedback stage, split into rewarded (win) and unrewarded (loss) trials. Significant clusters are identified using a permutation t-test between two conditions, with multiple comparisons corrected. (B) The ERP of the fictitious play update was calculated for significant electrode clusters, split into low, medium, and high terciles. Significant clusters are computed using a full-map GLM and corrected for multiple comparisons with the cluster permutation method. (C) ERP of the influence play update calculated for the significant clusters of electrodes split into low, medium, and high terciles. Significant clusters are computed using a full-map GLM and corrected for multiple comparisons with the cluster permutation method.

Previous research has shown that reward encoding and belief updates are represented in distinct brain regions, including the nucleus accumbens, TPJ, and dmPFC (Konovalov et al., 2021). Although we could not directly link our activations to specific brain areas, we could now examine whether updates in rewards and beliefs occur simultaneously or sequentially. We found that learning-update effects were temporally distinguishable from reward-outcome effects, with the fictitious update occurring earlier (Figure 2). Specifically, a whole-scalp sensor-level GLM correlating the amplitude signal and continuous measures of the model-specified learning updates revealed a negatively signed response to the fictitious play update (first-order belief, based on the opponent’s choice) within a short time window of about 100-300 ms, with the peak around 250 ms after feedback onset. The relevant cluster of electrodes was concentrated in the centroparietal region (Figure 2B). For the influence update, the analysis revealed a cluster that only partially overlapped with the fictitious play update, both in location and in timing: It was associated with activity in a separate set of electrodes in the centroparietal region, roughly between 250-500 ms after stimulus onset (Figure 2C).

These results provide evidence for the sequential nature of computational processes during social decisions. Given the sequential processing of these decision variables in the brain, our findings suggest it may be possible to selectively influence their processing across different contexts, for example, by adjusting feedback presentation and duration. This temporal separation may thus be crucial for understanding updates in environments with multiple feedback channels.

### Subjective values of options are reflected in the EEG signal

A key assumption of learning models is that the brain creates, updates, and retrieves subjective decision values. Previous studies have associated increased activity in the vmPFC and striatum with higher values of presented or selected options (Bartra et al., 2013), and activity in the anterior cingulate cortex (ACC) and dmPFC with decision conflict or differences in the values of options (Venkatraman et al., 2009; Venkatraman & Huettel, 2012). In a typical value-based task, the values shown on each trial are independent; here, we aimed to identify signals for values that are updated dynamically across trials. Furthermore, EEG enabled us to distinguish different aspects of information processing over time by analyzing signals locked to stimulus onset or response timing.

We assumed that the brain needs to compare the updated option values during the choice stage. Confirming this hypothesis, we did not find any activity significantly correlated with the (subsequent trial) value signals during the feedback stage (p > 0.05). Also consistent with our predictions, we found evidence for value encoding during the choice stage: Activity in a central cluster of electrodes was correlated with the continuous (signed) value difference between the two options within a temporal window around 250 ms after stimulus onset (Figure 3A). However, around the same time window (starting earlier, around 150 ms after stimulus onset), we also detected a frontal signal correlated with the (signed) influence update value from the previous trial (Figure 3B), suggesting a “recall” of the update that reflects the participant’s own choices. We found no activity significantly correlated with the fictitious play update (p > 0.05) and thus no evidence of a corresponding “recall” of the update reflecting the opponent’s previous choice.

**Figure 3.**
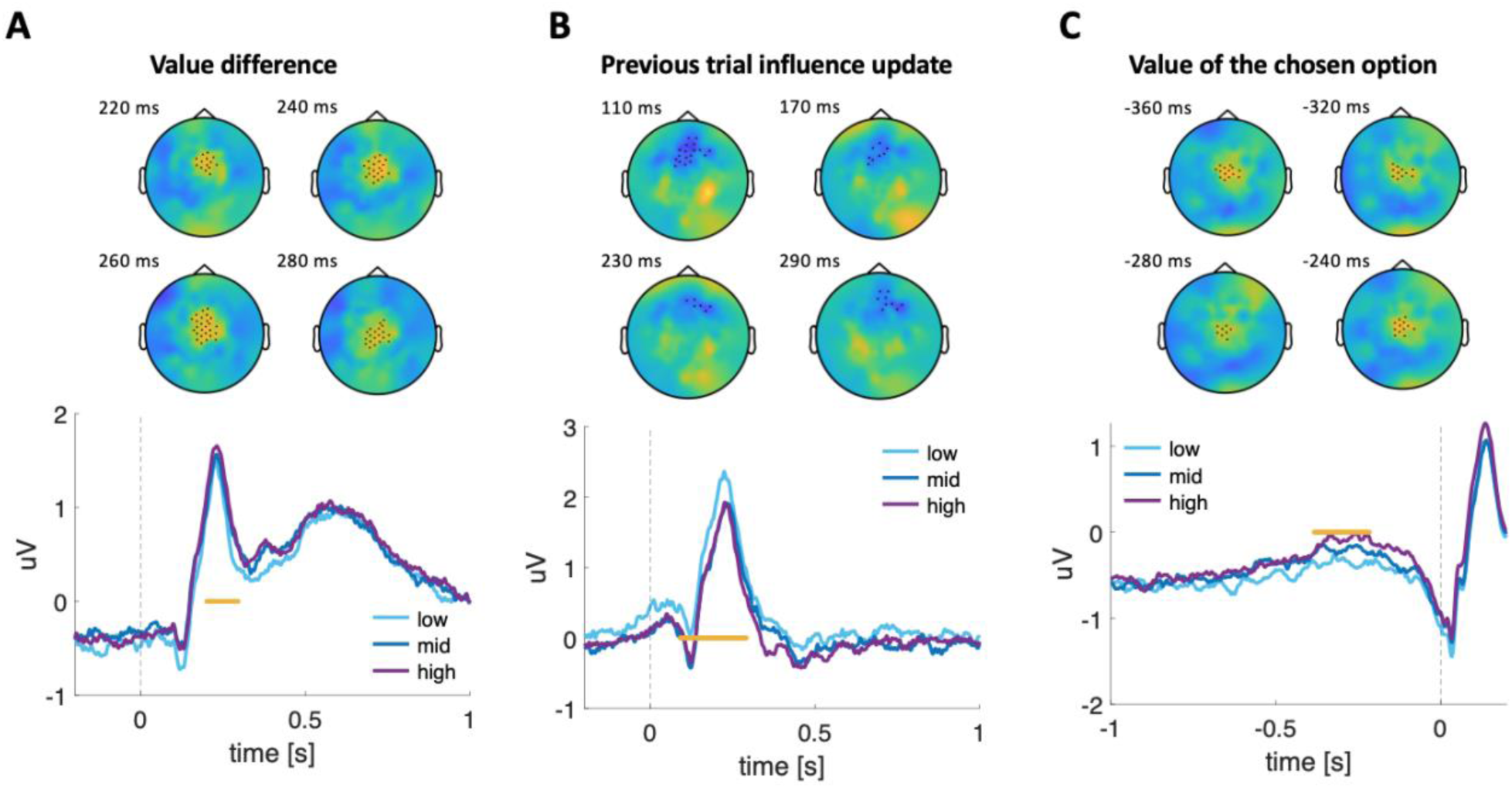
Choice stage EEG results. The topographical maps show the significant clusters (marked with black dots) at the indicated time after option onset, with yellow indicating positive correlation and blue indicating negative correlation. The ERPs below are calculated for the significant clusters, split into low, medium, and high terciles. Tercile splits were used only for visualization; statistical inference was performed on continuous trial-level regressors. Yellow lines in the ERPs reflect the (approximate) time of significance at p < 0.05. Significant clusters are computed using a full-map GLM and corrected for multiple comparisons with the cluster permutation method. (A) EEG signals related to the option value difference, locked to the stimulus onset (shown as 0). (B) EEG signals related to the previous trial influence update, locked to the stimulus onset. (C) EEG signals related to the value of the chosen option, locked to the response time (timepoint 0).

In the analysis using timings locked to the participant’s response, we did not detect any signals that correlated with value difference and influence update. However, we observed encoding of the chosen option’s value in a centrally located cluster of electrodes that showed an accumulation-to-bound pattern, which differentiated the value from approximately −400 to −200 ms before the response, but reached the same amplitude at the moment of response (Figure 3C). This activity pattern is typical of DDM-like processes (Hu et al., 2023; Polanía et al., 2014); in the next section, we examine this pattern in more detail.

### Parietal EEG signals reflect evidence accumulation for subjective values

Focusing on the response-locked analysis, we aimed to identify EEG signal patterns that correspond to the accumulation process in our DDM. Our model assumes that the integrated value signal (value difference) is transformed into a response during the final stage of the decision process. The DDM has been shown to describe this process as the dynamic accumulation of evidence (value difference) within the trial.

To describe and predict evidence-accumulation traces in the EEG data, we used a method previously employed in our lab (Hu et al., 2023; Polanía et al., 2014). Specifically, we used only half of the data (odd trials) and simulated the drift-diffusion process for each trial 100 times (see Methods). We then ran a regression on the raw amplitude signal to identify electrodes showing a significant correlation with the simulated millisecond-by-millisecond accumulation trace.

This identified one centroparietal cluster positively correlated with the accumulation trend and another centrofrontal cluster negatively correlated (Figure 4A, left panel). We then aimed to predict EEG activity out of sample using the remaining data (even trials). Following the previous literature on the centroparietal positive potential (CPP, see (Kelly & O’Connell, 2013)), we chose the cluster of 5 electrodes with the strongest positive correlation (Figure 4B, left panel) and predicted activity in this cluster during the even trials, employing the regression model trained on the odd trials.

**Figure 4.**
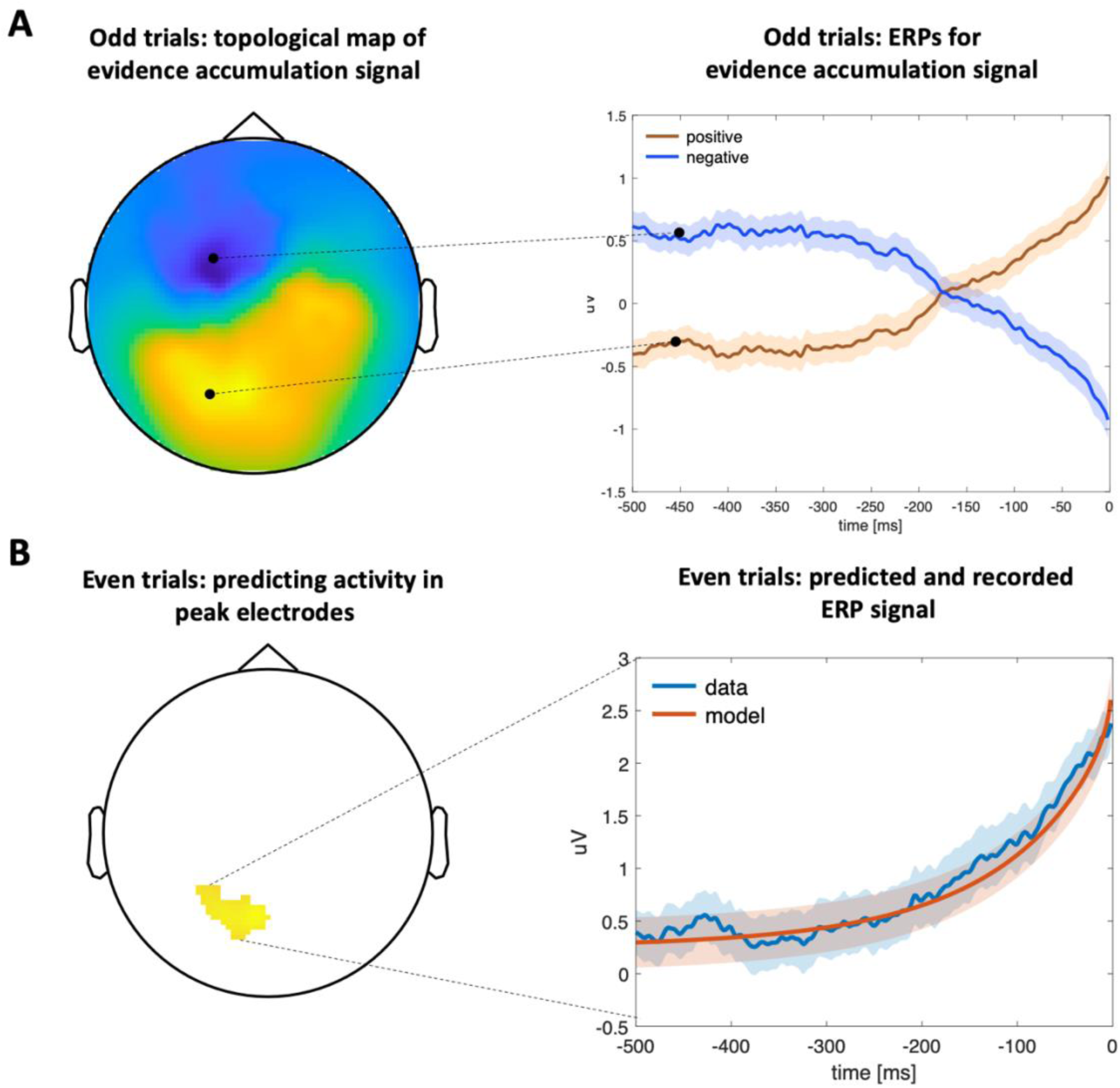
EEG: evidence accumulation traces. (A) Odd trials: significant clusters of EEG activity showing significant correlation (using a full-map GLM) with the trial-by-trial simulation of the DDM evidence accumulation traces (left panel), locked to the response time (shown as 0). ERPs are plotted for the electrodes in the positive and negative clusters showing a significant correlation with the simulated traces (right panel, p < 0.05, corrected for multiple comparisons). (B) Even trials: empirical (in blue) and predicted (in red) evidence accumulation traces (right panel) for 5 peak electrodes in the positive accumulation cluster (left panel).

We found that the EEG signal in this parietal cluster on even trials indeed followed the evidence-accumulation trajectories predicted by the accumulation model trained on odd trials (Figure 4B, right panel). This result aligns closely with findings in the value-based and perceptual domains, suggesting that EEG signals can reflect a domain-general, DDM-consistent accumulation process in the brain that shows similar patterns across different types of decisions (Hu et al., 2023; Kelly & O’Connell, 2013; Polanía et al., 2014). Although this signal may partly reflect amplitude changes associated with motor responses, the correlation between the trial-specific drift rate and the value of the chosen option (as shown in the previous section) strongly indicates that these accumulation processes are specifically linked to action-value computations rather than merely a simple ramp-up signal reflecting basic motor preparation.

### Centroparietal signal coherence reflects individual task performance

So far, our results suggest that the relative values of the options (H and T) are translated into actions during the choice process via a DDM-consistent evidence-accumulation mechanism. This indicates that value must be communicated from the value-computing regions to the areas involved in final integration and decision-making. Previous research has proposed that phase synchronization between regions facilitates efficient information transfer (Hu et al., 2023, 2026; Polanía et al., 2012, 2014, 2015; Siegel et al., 2012). EEG enabled us to examine the inter-regional gamma-band coherence of signals in electrode clusters associated with the chosen value and evidence accumulation (aligned to the response time). Prior studies have shown that such coherence is causally necessary for stable evidence accumulation (Hu et al., 2026; Polanía et al., 2014, 2015; Siegel et al., 2008). For this exploratory analysis, we investigated whether coherence between the two significant clusters identified in response-locked data (the central cluster related to the chosen option’s value and the parietal cluster linked to evidence accumulation processes) correlates with each individual’s performance level or other behavioral tendencies (Figure 5A).

**Figure 5.**
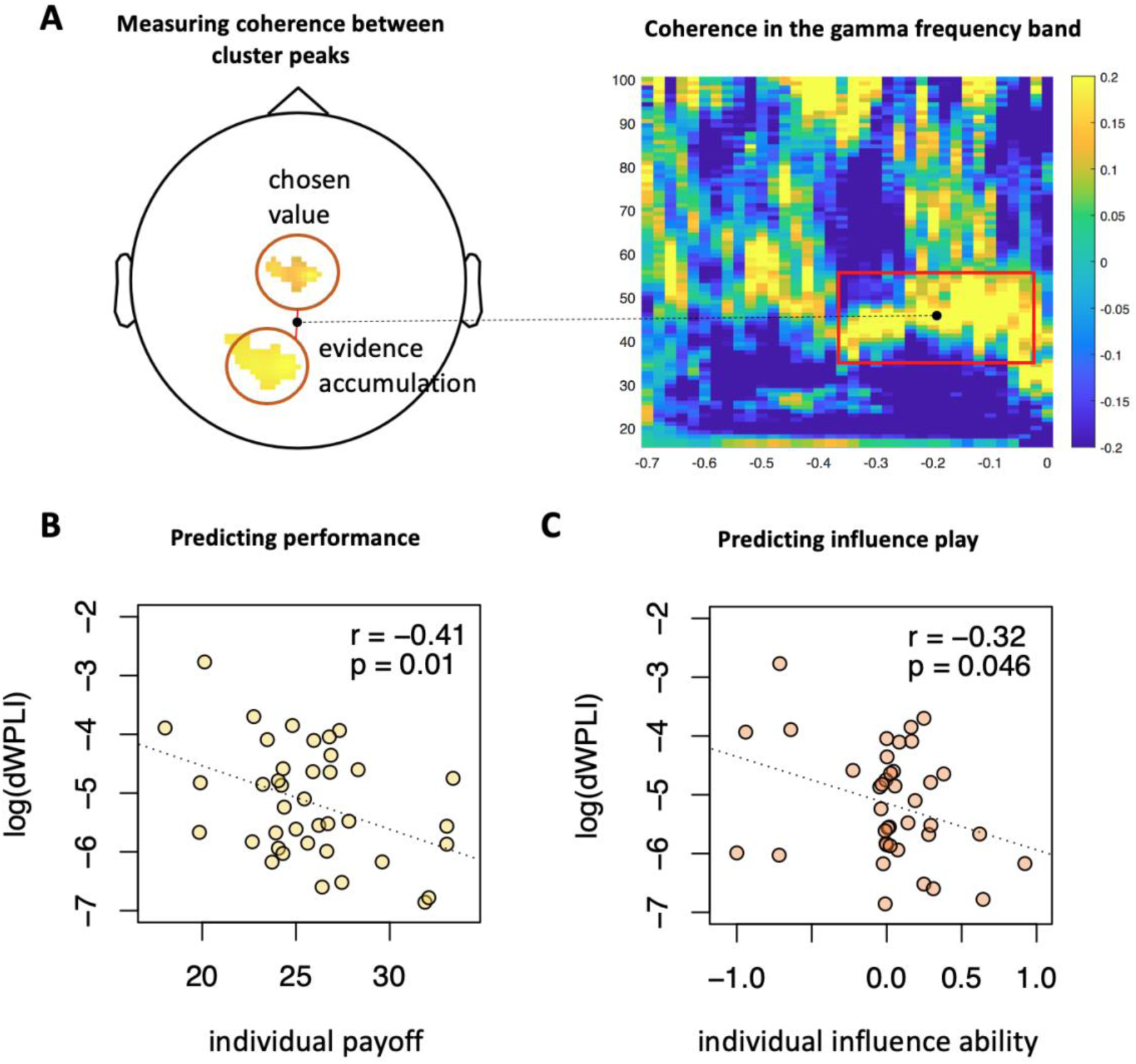
EEG: coherence between the central and parietal electrodes. (A) Left panel: peak electrodes identified in the response-locked analyses (chosen value and evidence accumulation). Right panel: gamma-band coherence between the two clusters, calculated as dWPLI. (B-C) Pearson correlation between the individual coherence measure (extracted from the time courses and frequencies shown in A) and performance (B) or influence play parameter 𝜅 (C).

We computed the response-locked analyses for each participant using the debiased weighted phase lag index (dWPLI, Vinck et al., 2011). Across all participants, we found a significant positive coherence band between 30 and 60 Hz within the time window of around 100 to 400 ms before response execution (Figure 5A), suggesting systematic functional communication between these two clusters just prior to choice.

We then calculated the coherence value within this significant cluster for each participant separately and correlated the individual values with the corresponding behavioral variables. This revealed a moderate yet significant negative correlation between the participant’s performance and their individual dWPLI (r = −0.41, t(37) = −2.7, p = 0.01, Figure 5B), as well as a negative correlation between the participant’s influence play parameter and the dWPLI (r = −0.32, t(37) = −2.1, p = 0.046, Figure 5C). There was no significant correlation between the dWPLI of the two selected clusters and the individual tendency to use fictitious play (r = 0.14, t(37) = 0.83, p = 0.41). We acknowledge that this analysis is somewhat underpowered and therefore consider these results exploratory, but note they partially align with previous research (Polanía et al., 2014, 2015). However, a notable difference is the sign of the correlation: in our study, higher coherence is associated with lower individual performance. This may reflect that the clusters used in our study and in previous studies were identified using different latent variables, which may serve somewhat different functions in the choice process. Alternatively, the negative correlation may indicate that participants with lower performance are more taxed by the task and therefore recruit stronger integration to select their actions. Regardless, our findings suggest that coherence between the central and parietal regions, which are involved in evidence accumulation and value computation, might also be essential for successful strategic decision-making.

## DISCUSSION

We used EEG to characterize the temporal dynamics of the decision-making processes underlying strategic choices, where choice options are fixed but decision values change over time. In our repeated strategic game, choices and response times were well captured by a computational approach that combines trial-by-trial updating of first-and second-order beliefs with a drift-diffusion process that accumulates the difference in subjective values between the two options available to the player. This provides direct evidence for models describing the underlying cognitive variables in strategic social decisions. Our EEG findings identify neural signatures corresponding to these model components, such as subjective values and belief updates, and offer new insights into the timing of these updates. Finally, our data reveal electrode clusters linked to the evidence accumulation process of the DDM and, consistent with previous research on individual decision-making, showed that coherence of these areas with value-sensitive regions relate to each participant’s strategic performance.

### Timing of belief updates

Our analysis of the feedback phase showed that belief updates (first- and second-order beliefs) were linked to similar spatial locations but were temporally structured differently: the fictitious-play update occurs first (peaking around 250 ms), and influence-update processing lasts up to 600 ms after feedback onset. Both update signals are correlated with activity in the centroparietal electrodes. Although we cannot localize this activity to specific cortical areas, previous fMRI literature suggests that these results could reflect activity in the dmPFC (for fictitious play (Hampton et al., 2008; Zhu et al., 2012)) and TPJ (for influence play (Hampton et al., 2008; Hill et al., 2017; Konovalov et al., 2021)). These sensor-level timing differences are consistent with the possibility that computations linked by prior fMRI work to dmPFC and TPJ unfold at different times.

One open question in learning and performance monitoring is whether outcome evaluation simply updates the choice values, with the actual choice computation occurring only when the options reappear and actions need to be selected, or if it instead creates a future action plan immediately upon feedback presentation (when the task structure permits). This question is motivated by RL models of feedback-related medial frontal signals and by extensive evidence that outcome-locked EEG components in the ∼200–600 ms range predict future behavioral adjustments (Sambrook & Goslin, 2015; Walsh & Anderson, 2012). For example, studies of trial-to-trial switching and reversals support the idea that feedback processing can shape future choice policies rather than just reflect retrospective evaluation (Chase et al., 2011; Cohen & Ranganath, 2007; Fischer & Ullsperger, 2013; Gehring & Willoughby, 2002; Holroyd & Coles, 2002; San Martín, 2012; San Martín et al., 2013). Computational modeling combined with eye-tracking has shown that, in some model-based learning scenarios, choices can be determined before the next trial begins, and the decision process need not align with the experimenter-defined choice screen (Konovalov & Krajbich, 2016). If value comparison for trial *t+1* had already occurred during feedback on trial *t*, we would expect feedback-locked signals of next-trial decision variables, such as the chosen value or the value difference, but we did not observe this. Instead, we found that the influence-update signal triggered by feedback was reactivated during the following choice period, supporting a two-stage account where feedback updates latent beliefs or values, and their comparison and commitment to action mainly happen once the options and motor mappings are set, consistent with evidence accumulation models and centroparietal decision signals (Frank et al., 2015; O’Connell et al., 2012; Pisauro et al., 2017).

### Value comparison and evidence accumulation

We identified a central cluster of electrodes that correlated with the action-value difference (using stimulus onset as the reference) and with the value of the chosen option (using response time as the reference). Along with the ERPs linked to win-vs-loss processing at outcome, these findings align with previous EEG research on reward, reward prediction error, and value processing, with this signal linked to activity in the vmPFC and nucleus accumbens (Cohen et al., 2007; Flores et al., 2015; Gorka et al., 2015; HajiHosseini & Holroyd, 2015; Hoy et al., 2021; Keren et al., 2018; Silvetti et al., 2014). Additionally, our framework suggests that stored values are converted into choices at the precise moment when an action is prompted, a process that is difficult to detect with fMRI. By aligning the signal to response onset, we observed that the signal amplitude in a central cluster of electrodes matched a DDM-like process, with the signal distinguishing the value of the chosen option approximately 300 ms before the response but reaching the same level at the moment of response (Figure 3C). These results are consistent with prior research on evidence accumulation in different contexts, in humans and primates (Donner et al., 2009; Gluth et al., 2012; Huk & Shadlen, 2005; Hunt et al., 2012; Kelly & O’Connell, 2013; Kiani et al., 2008; O’Connell et al., 2012; Pisauro et al., 2017; Polanía et al., 2014; Roitman & Shadlen, 2002; Shadlen & Newsome, 2001), but extend this research to the domain of strategic decision-making.

Thus, our results indicate that a common process underlies many different types of value-based decision-making, including dynamic social decisions. Previous studies have shown that the DDM and other evidence accumulation models can serve as decision rules for simple reinforcement learning situations (Fontanesi, Gluth, et al., 2019; Fontanesi, Palminteri, et al., 2019; Miletić et al., 2020; Pedersen et al., 2017), with EEG data supporting this mechanism (Bridwell et al., 2018; Frank et al., 2015). Here, we extend these findings to more complex social decisions, suggesting possible future research on the further separation of belief components (beliefs, values, and preferences) in strategic decisions, both temporally and topographically (Konovalov, 2026).

### Individual behavior linked to gamma-band coherence

Previous research indicated that frontoparietal connectivity might be a crucial component of the evidence accumulation process, facilitating information sharing between value-encoding and decision-making regions (Basten et al., 2010; Philiastides et al., 2010; Pisauro et al., 2017). Specifically, changes in gamma-frequency band coherence are associated with shifts in performance during perceptual and value-based decision-making (Gregoriou et al., 2009; Vinck et al., 2013; Womelsdorf et al., 2006), with modulation of phase coupling between frontal and parietal areas increasing decision accuracy (Polanía et al., 2015). Our results align with these findings, demonstrating that regions involved in evidence accumulation and value encoding exhibit significant changes in coherence during strategic decisions, with coherence strength linked to individual performance and the chosen influence strategy. These findings are consistent with functional coupling between the selected electrode clusters during response preparation, but do not establish directionality or causal information transfer and require further research. These findings suggest that frontoparietal (or centroparietal) coherence could serve as a key common mechanism underlying value-based decisions, a hypothesis that could be explored using causal stimulation techniques (Hu et al., 2026; Polanía et al., 2015; Soutschek et al., 2021).

To summarize, our findings provide important validation of both learning and decision models of strategic behavior, supporting the idea that the brain stores and updates action values during social decisions, uses feedback to revise beliefs, and engages in dynamic comparison during choice. These findings can provide new insights into learning and decision-making, helping break down complex behaviors into fundamental computational components.

## METHODS

### Participants

We recruited 84 participants for the study; 42 took part in the EEG part, while the other 42 served as their opponents in a different room. All EEG participants (17 females, mean age: 25, SD = 2.7) were right-handed, free of neurological disorders, had normal or corrected-to-normal vision, and were medication-free. Participants in the EEG study received between 67 and 180 CHF, and the opponents received between 48 and 93 CHF for their participation, based on their performance in the task (1 CHF = 1 USD at the time of the experiment). Two participant pairs were excluded from the behavioral analyses due to a data encoding error, and one participant was excluded from the EEG analysis due to excessive EEG artifacts. The data from the remaining 39 participants were used for EEG analyses. All participants provided written consent, and the study was approved by the ethics committee of the Canton of Zurich.

### Task

Each EEG participant played a standard strategic game, the Inspection game (Hampton et al., 2008; Hill et al., 2017), against a real human opponent (another participant). We removed the typical labels of the players and actions, such as “employee/employer” and “work/shirk,” to avoid cultural connotations. Participants were instructed to choose between two options (labeled H and T to minimize visual stimulation), while their opponent’s choice was presented as UP or DOWN. If the participant chose H and the opponent chose UP, or the participant chose T and the opponent chose DOWN, the participant earned 50 points, while the opponent received 0 points; the opponent scored 100 points if the participant chose H and the opponent chose DOWN, and 25 points if the participant chose T and the opponent chose UP, with the participant receiving 0 points in both cases. Thus, the payoff matrix was identical to that of the Inspection game (see Figure 1).

Both the EEG participants and their opponent made their choices simultaneously. Here, we will describe the EEG participant’s setup; the opponent’s interface was identical. In each trial, a fixation cross was displayed on the screen, with placeholders (“X”) for the payoffs and choice options (Figure 1). Participants could rest between trials until they initiated the next trial by fixating the central cross for 0.5–1 second (controlled by an eye-tracker and uniformly jittered). The interface then displayed two options on either side of the fixation cross: H and T. We randomized the positions of the options (left/right) on each trial to prevent participants from preparing their motor response in advance. Participants used a standard EEG-compatible button response box to select the left option (with their index finger) or the right option (with their middle finger). Participants had 3 seconds to respond; after a button press, the chosen option was highlighted in blue (all colors were approximately luminance-matched). 3.5 seconds after the options appeared, the opponent’s choice was displayed on the screen. The top row of payoffs indicated UP, and the bottom row indicated DOWN. The relevant payoff (0 or 50 points) was highlighted in blue. Feedback remained on the screen for 1.5 seconds before a new trial began.

The opponent played in a different room using a standard keyboard, without eye tracking, while the computers were connected over the LAN. The game was presented similarly (as a choice between H and T), but the opponent saw their own payoffs on the screen.

The participants played the game for eight blocks, each consisting of 50 trials followed by a short rest period. The two participants arrived in separate rooms and did not communicate to avoid prior beliefs and non-strategic behavior; we ensured that both understood they were playing against a real opponent. Participants practiced the task in 10 unpaid trials. At the end of the experiment, we converted the total points earned in the task at a rate of 150 points = 1 CHF and added a 20 CHF show-up fee.

### EEG recordings

All testing took place in the SNS Lab at the University Hospital Zurich, in a soundproof and electromagnetically shielded recording chamber. We recorded EEG signals using 128 scalp sites with sintered Ag/AgCl electrodes mounted in an equidistant hexagonal layout in a Waveguard Duke 128-channel cap (http://www.ant-neuro.com/).

We connected the cap to a 128-channel QuickAmp system (Brain Products, Munich, Germany). We referenced the EEG signals online to the average reference. We kept the impedance below 5 kΩ for all the electrodes throughout the experiment. We amplified the signals using a bandpass filter from 0.016 to 100 Hz and digitized them online at a sampling frequency of 500 Hz. We set up the cap on the participant’s head before they entered the testing chamber.

### Eye tracking

We used an SR Research EyeLink-1000 (http://www.sr-research.com/) eye tracker to record participants’ fixation patterns at a sampling frequency of 500 Hz. Before each trial, we instructed the participants not to blink and to keep their eyes fixated at the central fixation cross for 0.5-1 second before the trial started.

### Computational model

To fit the participants’ data, we used a computational model that combines a strategic learning model established in the literature, known as the influence model (Hampton et al., 2008; Hill et al., 2017), and a standard drift-diffusion choice model (Ratcliff & McKoon, 2008) with fixed bounds and a fixed starting point (0.5), where the drift rate depends on the difference in values of the choice options (H and T).

Specifically, the model assumed that participants updated their beliefs about the opponent’s action on each trial after receiving feedback (the opponent’s action, UP or DOWN):

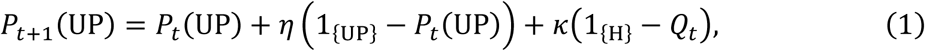

where 𝑃_𝑡+1_(𝑈𝑃) is the subjective probability (belief) that the opponent will play UP on the subsequent trial; 𝑃_𝑡_(UP) is the same (cached) belief for the current trial; 1_{UP}_ is an indicator function that is equal to 1 if the opponent played UP on the current trial and 0 otherwise; 1_{H}_ is the indicator for the participant’s choice on the current trial (= 1 if H, 0 otherwise); 𝜂 is the participant-level parameter that reflects the strength of the first-order (fictitious play) belief update; 𝜅 is the participant-level parameter that reflects the strength of the second-order (influence) belief update. 𝑃_𝑡_ is truncated to [0,1] to keep the beliefs within the probability measure.

𝑄_𝑡_ is the second-order belief computed as an inverse of the opponent’s assumed DDM-based choice rule:

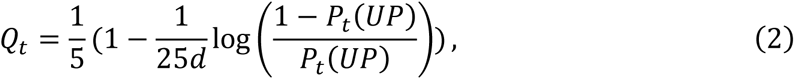

where 𝑑 is the participant-level parameter (drift rate scaling). 𝑄_𝑡_ is truncated to [0,1] to keep the beliefs within the probability measure. For parsimony, we used the same scaling parameter 𝑑 for the participant’s drift rate and for the inverse choice function used to infer second-order beliefs.

The updated subjective beliefs are used to compute the values of the choice options on the next trial, computed as expected values of each choice (H or T):

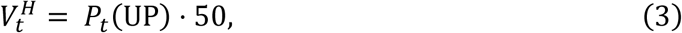

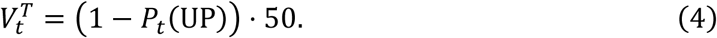

The values of the options are used as an input for the drift rate of the DDM model:

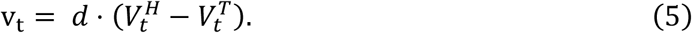

We then used the drift rate 𝑣_𝑡_, a boundary separation parameter 𝑎, and a non-decision time parameter 𝜏 to compute the likelihood of the RT and choice (H or T) on each trial using the RWiener package (Wabersich & Vandekerckhove, 2014), and maximize the total likelihood over all trials to obtain the individually fitted parameters 𝑑, 𝑎, 𝜏, 𝜂, and 𝜅. We searched over a random grid of 250,000 parameter sets and used the 10 sets with the highest likelihood as starting points for gradient optimization using *optim* in R; the final values were selected from the best-fitting set among these 10.

For the optimal set of parameters, we calculated the following values for the EEG analyses: 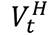 and 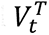 as values of the options on the current trial, (1_{UP}_ − 𝑃_t_ (UP)) as the fictitious play update on trial 𝑡, (1_{H}_ − 𝑄_𝑡_) as the influence update on trial 𝑡. For the posterior predictive checks, we simulated the model 100 times for each trial (Figure 1).

We compared this model to several competing models: Q-learning, fictitious play (assuming 𝜅 = 0), and a model where the difference in values modulates the starting point of the DDM process rather than the drift rate; these alternative models provided a worse fit to the data (Supplementary Figure 1). We also compared this model to a standard influence model with a softmax choice function. DDM-based models were compared using the joint choice–RT likelihood and BIC. The DDM model was compared with the softmax influence model only on choice-prediction accuracy as their likelihoods condition on different data. Both models provided similar mean prediction accuracy (Supplementary Figure 1).

### EEG preprocessing

We preprocessed the EEG data using Fieldtrip (Oostenveld et al., 2011) implemented in Matlab R2020b (MathWorks). We removed eye movements and other noise artifacts using independent component analysis, selecting components based on their topography and power spectra. We used a discrete Fourier transform to filter line noise.

For the analysis of choice onset (H and T presented on the screen), we extracted data from a time window from −200 ms to +1000 ms relative to stimulus onset. For the response-locked analysis, we extracted the data from the -1000 to +200 ms window around the button press. For the feedback analyses, we extracted the signal from the −200 to +1000 ms window relative to the onset of the opponent’s choice (i.e., the feedback). We visually inspected all individual trials, removing those with high amplitude variance (e.g. muscle artifacts) for the EEG analyses (Chen et al., 2019). Overall, we rejected an average of 8.7% ± 5% trials per participant.

### EEG analysis

For the analysis of the representation of model-based variables (value difference, chosen value, fictitious and influence updates) in the EEG signals, we ran a full-map GLM regressing the EEG amplitude in each timepoint of the window of interest (-0.2 to 1 s for the choice and feedback-locked analyses, -1 to 0.2 s for the response-locked analysis) on the variable of interest in each trial of the task and each channel, for each participant separately. We then used Fieldtrip’s permutation analysis of the regression betas (measuring parametric effect strength) to identify significant clusters of activity, corrected for multiple comparisons at p(FWE) < 0.05. To plot the ERPs, we extracted EEG amplitudes from all electrodes identified as significant in these analyses and split them into three terciles based on the underlying model-based variable.

To account for multiple comparisons, we identified clusters using two-sided *t* statistics (*p* < 0.05 at the channel level) and required at least three significant neighboring channels within each cluster. We defined the cluster size as the integral of the t-scores across all channels in the cluster. The significance was tested with a permutation statistic (performed 5,000 times with shuffled labels to generate a distribution of cluster sizes under the null hypothesis) (Maris, 2012). We applied this threshold to all the whole-brain EEG analyses.

### Evidence accumulation EEG analysis

To identify the EEG signals associated with the evidence accumulation process, we employed an approach previously established in our lab (Hu et al., 2023; Polanía et al., 2014). We simulated the DDM model 100 times per trial and averaged the resulting curves to obtain a trial-level predicted accumulation trace. Next, we used data from odd trials to identify the channels most closely related to the predicted evidence-accumulation traces. Specifically, for each channel and trial within each participant, we analyzed a time window from -500 ms to the response by regressing EEG amplitude on the predicted evidence value at each time point. We then applied the resulting regression model and, using the top five channels from the positive cluster (as selected by regression beta values), predicted the EEG signal in the independent even trials for the same set of channels.

### EEG connectivity analysis

We used the debiased weighted phase lag index (dWPLI) (Vinck et al., 2011) for the connectivity analyses. This statistic is based on the imaginary component of the cross-spectrum, ensuring that differences in power do not affect coherence measurements and eliminating small-sample bias when estimating the phase lag.

We calculated the spectral estimates (power and cross-spectral density) for each response-locked time window using the multitaper method implemented in FieldTrip. The time-frequency analyses were performed within the 16-100 Hz frequency band. We set the temporal sliding window length to 8 cycles, with a step size of 0.02 s. The frequency smoothing width was set to 0.3f, with a frequency resolution of 1 Hz steps. Next, we calculated the dWPLI for all potential pairwise connections between the selected peak electrodes and the evidence accumulation peak electrodes (Figure 5A). Our focus was on signals in the gamma band (30-90 Hz) within 500 ms prior to responses. Finally, we extracted the average dWPLI measure for each participant from the time-frequency cluster identified as significant across all participants, using permutation methods and correcting for multiple comparisons at p < 0.05.

## Acknowledgments

We thank Gábor Stefanics for helping with setting up the EEG system. This project has received funding from the European Research Council (ERC) under the European Union’s Horizon 2020 research and innovation programme (grant agreement No 725355).

## Author contributions

A.K., J.H., and C.C.R. designed the research and hypotheses. A.K. and J.H. collected the data. A.K. and J.H. analyzed the data. A.K., J.H., and C.C.R. co-wrote the paper. C.C.R. supervised the project.

