## Supplementary materials for "Neural Dynamics of Belief and Value Computations Guiding Strategic Social Decisions"

**SUPPLEMENTARY FIGURES**

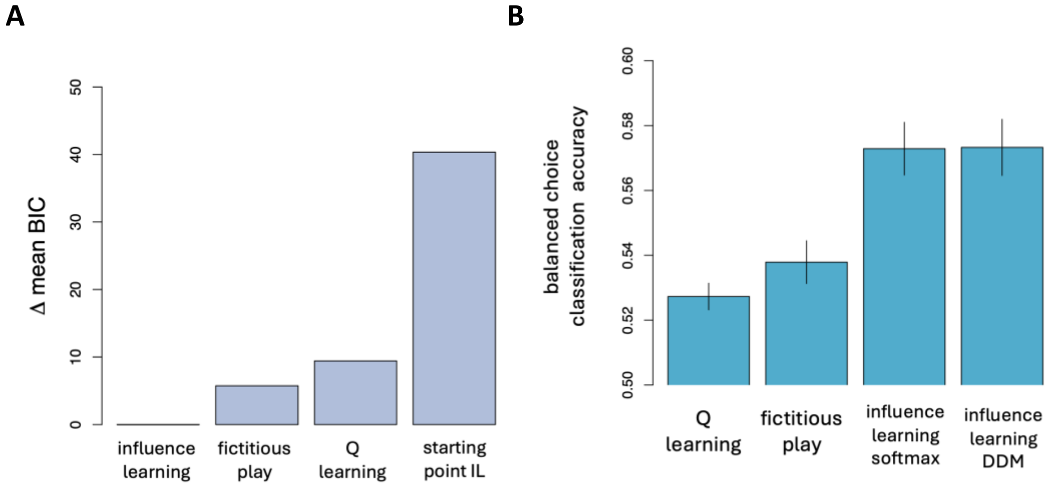


**Supplementary Figure 1.** (A) Difference in mean participant-level Bayesian Information Criterion between competing computational models, including fictitious play, Q-learning, and the influence model with the starting point defined by the difference in action values. (B) Balanced choice prediction accuracy for the fitted models. The difference between the influence learning model estimated with the standard softmax choice function and the DDM choice function is not statistically significant.
